# BAP1 deletion in hepatocytes primes an inflammatory transcriptional response

**DOI:** 10.1101/2025.10.31.685807

**Authors:** William C. Nenad, Peyton Kuhlers, Ian R. Sturgill, Irene Biju, Max Bucklan, Lindsey Hernandez, Lee-Ching Zhu, Katherine A. Hoadley, Jesse R. Raab

**Author notes:** These authors contributed equally to the manuscript (William C. Nenad, Peyton Kuhlers). Corresponding Authors **Address for Correspondence:** Jesse R. Raab, PhD, Department of Genetics, University of North Carolina at Chapel Hill, 120 Mason Farm Rd., Chapel Hill, NC 27599.

## Abstract

**Background:** BRCA1-associated protein 1 (BAP1) is a deubiquitinase, frequently altered in cancers including hepatocellular carcinoma and cholangiocarcinoma. While Bap1 has been shown to play key roles in metabolism, maintenance of tissue homeostasis, and immune cell development, little is known about its normal functions in the liver in vivo. This study aims to identify Bap1 specific effects on the liver’s immune microenvironment and biological functions.

**Methods:** Using AAV8-mediated CRISPR/CAS9 genome editing we generated a mouse hepatocyte-specific model of Bap1 knockout to define the changes that occur in liver biology in an *in vivo* system and characterize how loss of Bap1 alters the liver’s response to injury. Single-cell resolution spatial transcriptomics were performed in conjunction with immunohistochemistry to analyze cell-type composition and immune cell recruitment changes. Bulk RNA-sequencing was performed for further assessment of the impact of Bap1 loss on transcription.

**Results:** Hepatocyte-specific depletion of Bap1 induced transcriptional changes shared with acute injury. We observed a strong dysregulation of inflammatory pathways associated with BAP1 loss. Moreover, the transcriptional response of Bap1 depletion in hepatocytes to damage was markedly different than in control liver, with Bap1-deleted livers showing a decreased hepatocyte identity based on gene expression. Spatial transcriptomics and quantitative texture analysis of immunohistochemistry revealed an altered immune environment prior to damage and an impaired recruitment of immune cells in Bap1 depleted livers after damage.

**Conclusions:** **Using a** hepatocyte-specific Bap1 deletion we identified Bap1 as a critical modulator in the liver’s immune cell response. We show that Bap1 loss leads to an inflammatory environment prior to damage and disrupts the recruitment immune cells. Our quantitative spatial analysis highlights the power of such approaches to characterize the spatial distribution of different cell types in a tissue.

## Introduction

Bap1 is a tumor suppressor gene encoding a deubiquitinase whose main function is to remove ubiquitination from Histone H2A at Lysine 119 (H2AK119ub). H2AK119ub is deposited by Polycomb repressive complex 1 (PRC1), making BAP1 a critical brake on transcriptional repression. Chromatin dynamics play a critical role in acute and chronic injury responses as well as liver cancer^1–4^.

H2AK119ub serves as a docking site for Polycomb repressive complex 2 (PRC2), where PRC1/2 dynamics have been shown to control hepatocyte maturation and fibrogenesis^5–7^. BAP1 inactivation is frequently observed in cholangiocarcinoma and a subset of hepatocellular carcinomas^8–10^. BAP1 loss in a hepatocellular carcinoma cohort from TCGA molecularly resembled cholangiocarcinoma, suggesting that BAP1 may regulate cellular plasticity in the liver^11^.

Bap1 is a key regulator in the liver, contributing to the maintenance of tissue homeostasis, normal liver function, and metabolism^12^. Artegiani et al. generated a human liver organoid model of BAP1 knockout and showed that BAP1 loss resulted in dysregulation of the epithelial cell state and identity^13^. BAP1 plays a central role in immune cell development and responses through its ubiquitination activity in B-cell, T-cell, and macrophage development^14–16^. Additionally, BAP1 loss in tumor cells has been shown to alter the tumor immune microenvironment^17–19^.

Despite these insights, the precise role of BAP1 *in vivo* in the liver and during the initial liver damage response is unknown. Given BAP1’s association with cancer and the importance of PRC1/2 in fibrosis, uncovering the function of Bap1 early in tissue injury is critical for understanding how its loss contributes to later disease phenotypes. Previous work used the combined deletion of Bap1 and Tp53 loss to study cell fate changes in organoids but have not explored the role of BAP1 *in vivo*^13^.

Here, we generate a mouse hepatocyte-specific model of Bap1 loss to explore and characterize the changes that occur in liver biology in an *in vivo* system. Our work demonstrates that hepatocyte-specific deletion of Bap1 induces widespread changes to the immune environment and liver damage response during acute injury. Using high resolution spatial analysis, we demonstrate changes to macrophage localization. Moreover, we show that Bap1 loss in the absence of damage induces an inflammatory response in the liver that is similar to the immune activation induced by damage.

Together, these data suggest a complex interaction of all cells in the liver tissue milieu that are impacted by Bap1 loss in hepatocytes, including important immune cell populations that influence liver health, homeostasis, and tumorigenesis.

## Methods

### Mouse Model

Cas9 Lox-Stop-Lox mice were purchased from Jackson Laboratory and bred in house (Strain #024857). 16-20 week old female mice were injected via the lateral tail vein with 1e12 viral particles of AAV-sgRNA-TBG-Cre. At day 10 mice were injected with either 1:9 dilution of CCl4 in olive oil or vehicle (olive oil) intraperitoneally at 10uL/g bodyweight. 48 hours after treatment the left lobe was harvested for paraffin embedding and IHC and the right lobe was isolated for RNA. All animal experiments were approved by the Institutional Animal Care and Use Committee at The University of North Carolina at Chapel Hill

### Immunohistochemistry

Livers from animals were excised and the left lobe was fixed in 10% neutral buffered formalin for 24 hours at RT before paraffin embedding, slide sectioning, and H&E staining (UNC Pathology Services Core). For immunohistochemistry experiments, 5μm liver sections were deparaffinized by incubating 3 times for 5 minutes at RT in Citrisolv and rehydrated by two sequential incubations for 10 minutes in 100% Ethanol, 95% Ethanol, and then followed by 2, 5 minute washes in dH20. Antigen retrieval was performed in ImmunoRetriever Citrate unmasking solution in a Tintoretriever (Bio SB) on high pressure for 15 minutes. Sections were washed 3 times for 5 minutes in dH_2_O followed by incubating for 10 minutes in 3% hydrogen peroxide and finally washed in TBS with 0.1% Tween (TBST). Slides were incubated for 1 hour at RT in TBST + 5% Goat Serum for blocking. Primary antibodies were diluted in TBST + 5% goat serum (Cre 1:100 Cell Signaling, catalogue 15036S Lot #2, F4/80 1:500 Cell Signaling catalogue 70076 Lot 9) and incubated at 4 degrees overnight in a humidified chamber. Slides were then washed 3 times for 5 Minutes with TBST before addition of 1-3 drops of Signal Stain Boost Detection Reagent (Cell Signaling Technologies) directly to the slides and incubated for 30 minutes at RT. Slides were then washed 3 times for 5 minutes with TBST and 1 drop of Signal Stain DAB was diluted in 1mL Signal Stain Diluent before adding to tissue for 5 minutes. Slides were immersed in H_2_O for 5 minutes twice and counterstained with hematoxylin for 1-2 minutes and then soaked in Tap water for 1-2 minutes. Slides were then washed twice for 5 minutes in dH_2_O and then dehydrated by sequential incubates in 95% Ethanol, 100% Ethanol and Citrisolv before mounting with Signal Stain Mounting Media. Slides were imaged on a Leica Dmi8 or an Olympus VS200 slide scanner.

### IHC Quantification

Cre quantification using IHC slides was done in Qupath (version 0.5.0)^20^. Tiles were created to cover tissue and tiles that encompassed only tissue were selected for. Using these tiles positive cell detection was performed using the optical density sum in the Qupath module. The percent positivity was then graphically represented.

### H-Score F4/80 Quantification

F4/80 quantification using IHC slides was done in Qupath (version 0.5.0). 6 regions across each tissue were selected with a combined total area of ~2587212 um^2 ^20^. Built in positive cell identification was then performed using optical density (OD) sum and OD mean thresholds at .2,.4, and .6. H-scores were then computed per region and graphically represented by score using ggplot2 (version 4.0.0) ^21^. Anova was used to compute statistical significance between groups.

### qPCR

RNA was harvested with Monarch Total RNA extraction kit (NEB) and 1ug of RNA was converted to cDNA using Lunscript RT by 25 degrees for 2 minutes, 55 degrees for 10 minutes, and 95 fo1 minute. 2uL of a 1:10 dilution of cDNA was then used as input for qPCR with SssFast (BioRad) with 300nM gene-specific primers. Quantitation was performed as delta delta ct relative to Hprt and non-targeting control (Supplemental Table 1). BAP1 Forward GACCTTCAGAGTAAATGCCAGG, BAP1 Reverse ACCAACGTAGAAACCTTGCG, Tbp Forward GGGAGAATCATGGACCAGAA, TBP Reverse CCGTAAGGCATCATTGGACT.

### Histopathology Review

All samples were examined using hematoxylin and eosin (H&E) stained tissue sections. Slides were evaluated by a liver pathologist for zone 3 necrosis and steatosis. Samples were graded using a semi-quantitative scale. Zone 3 necrosis was graded by the varying severities of injury and extent of involvement in zone 3. Steatosis was graded by degree of swollenness and injury (Supplemental Table 2).

### RNA-Sequencing Analysis

RNA was extracted from fresh frozen liver samples using Applied Biosystems MagMAX mirVana Total RNA Isolation Kit (A27828). RNA sequencing libraries were generated from 500-1000 ng input RNA using Illumina TruSEQ Stranded mRNA kit (Cat 20020594). Library concentration was calculated using Qubit 1x dsDNA HS Array. RNA sequencing libraries were sequenced 2×50 on an Illumina Nextseq200. Reads were aligned to the reference genome mm10 (GRCm38_p6) using STAR 2.7.6a and genes were quantified ^22^with Salmon 1.4.0 to gencode vM25 ^23^. Quality control was performed using samtools sort 1.10, Picard 2.22.4 CollectRnaSeqMetrics, calculate max read length, flagstat, and run FastQC 0.11.9 ^24,25^.

Differential expression was performed with DESeq2 (version 1.42.1) using estimated counts from Salmon ^23,26^, Following differential expression, shrinkage was applied to each coefficient using the lfcShrink function with type = “apeglm” (version 1.24.0)^27^ (Supplemental Table 3). Results were visualized using ComplexHeatmap (version 2.18.0) and volcano plots with ggplot2 ^21,28^.

Interaction between genotype and treatment were estimated with DESeq2 using the following design: “~ treatment + genotype + treatment:genotype” ^26^. Interaction terms were shrunk and genes with significant coefficients (adjust p-value < 0.05) were used for further analysis. Log normalized expression values were median centered across samples and clustered using 1-pearson correlation distance and average linkage with the R package hclust (Supplemental Table 5).

### Gene Set Enrichment Analysis (GSEA)

Shrunken log fold changes were used for downstream gene set enrichment analysis. The package msigdbr (version 7.5.1) was used to extract c5 and c8 gene sets from bulk data using the *GSEA* function from clusterProfiler (version 4.10.1) ^29–34^. For enrichment/overrepresentation analyses, the *enricher* function was used. Gene sets were filtered by significance with a p-value threshold of 0.05, unless otherwise noted. Terms were grouped into “immune” and “non-immune” gene sets by keywords in (Supplemental Table 6). These were visualized as waterfall plots using ggplot2 ^21^.

### Bap1 Activity Score

BAP1 activity scores were calculated as a weighted sum as previously described ^8^. Briefly, BAP1-‘altered’ gene sets were identified by Sturgill et al. and the log2 normalized expression of the upregulated genes from this study were multiplied by +1 and downregulated genes were multiplied by −1 and summed to derive a final score (Supplemental Table 7). Spatial Transcriptomics Spatial transcriptomics data were generated using the Molecular Cartography platform (Resolve Biosciences). The technology is derived from single-molecule fluorescent *in situ* hybridization (smFISH) and produces an approximate spatial resolution of 300nm. Livers were harvested and immediately placed in O.C.T (TissueTek) before freezing in a dry ice/isopentane bath. 10μm fresh-frozen liver tissue sections section on a cryotome and placed on a Resolve Biosciences Molecular Cartography slide. Samples were shipped to Resolve Biosciences for processing and data generation.

A set of 100 targeted RNA probes were measured (Supplemental Table 8). Data was processed in the Resolve Biosciences online pipeline. Cell segmentation was achieved using Baysor and sample DAPI staining as a prior ^35^.

Putative cell types were assigned using robust cell type decomposition (RCTD) from the spacexr package (version 2.2.0) ^36^. The RCTD single cell reference was built from liver cell type annotations and expression data from the mouse steady state (StSt) liver cell atlas provided by Guilliams et al ^37^. Spatial cell types were called using default settings per-replicate by RCTD. Differences in cell type proportion across treatment conditions were determined using propeller (version 1.2.0)^38^.

### Image Analysis

F4/80 IHC slides were imported into R using OpenImageR (version 1.3.0) ^39^. Images were converted to greyscale. For each image, a patch of 200×200 pixels of background were selected for background intensity normalization. The pixel intensity of each image was divided by the mean of its background pixels. A 1500×1500 pixel area was selected for analysis (Supplemental Data Table 9). Images were tiled into 8×8 grids (187.5px x 187.5px) and filtered to remove tiles that encompassed mostly empty or veinous regions, using a gaussian blurring effect from EBImage (version 4.48.0) and a threshold of more than 10% of pixels in a tile in top 10% of pixel values of 1500×1500 pixel image ^40^. Each tile was rasterized using raster (version 3.6-31) and aggregated by taking a 3 × 3 pixel average using the aggregate function from terra (version 1.8-15) ^41,42^. Filtered and aggregated tiles were run in Grey-level Co-Occurrence Matrices (glcm) algorithm (version 1.6.5) using contrast, homogeneity, entropy, second moment, variance, mean and dissimilarity statistics across 0°, 45°, 90°, and 135° shifts with the number of grey levels (n_grey) set to 4^43,44^. For each tile, the mean, standard deviation, quantile 1 and quantile 3, kurtosis, and skewness were calculated for each glcm metric with e1071 package (version 1.7-16) and used in prcomp (stats-package) for PCA analysis ^45^. Tiles were visualized on the PCA using ggimage (version 0.3.4) ^46^.

## Results

### BAP1 loss alters immune function

We evaluated the role of Bap1 in the liver and response to damage by generating a hepatocyte-specific mouse model of Bap1 loss by injecting CAS9-LoxStopLox mice with AAV8 encoding Cre recombinase under the hepatocyte specific Tbg promoter and a guide RNA targeting BAP1 or a non-targeting control, herein referred to as sgBAP1. After 10 days, non-targeting control (NTC) and sgBap1 mice were treated with either carbon tetrachloride (CCl_4_) or vehicle and 48 hours later livers were harvested (Figure 1A). We observed robust IHC staining for Cre expression and transfection efficiency was comparable across all experimental groups with overall mean Cre positivity of 78.1% (ANOVA p = 0.327, Supplemental Figure 1A,B). Reduction of Bap1 RNA expression in sgBap1 livers was confirmed by both qPCR and bulk RNA-seq (t-test p=9e-5 and Veh: NTC vs sgBap1 = 2.89 e-7, CCl_4_: NTC vs sgBap1 = 5.73 e-6, respectively, Supplemental Figure 1C,D).

**Figure 1.**
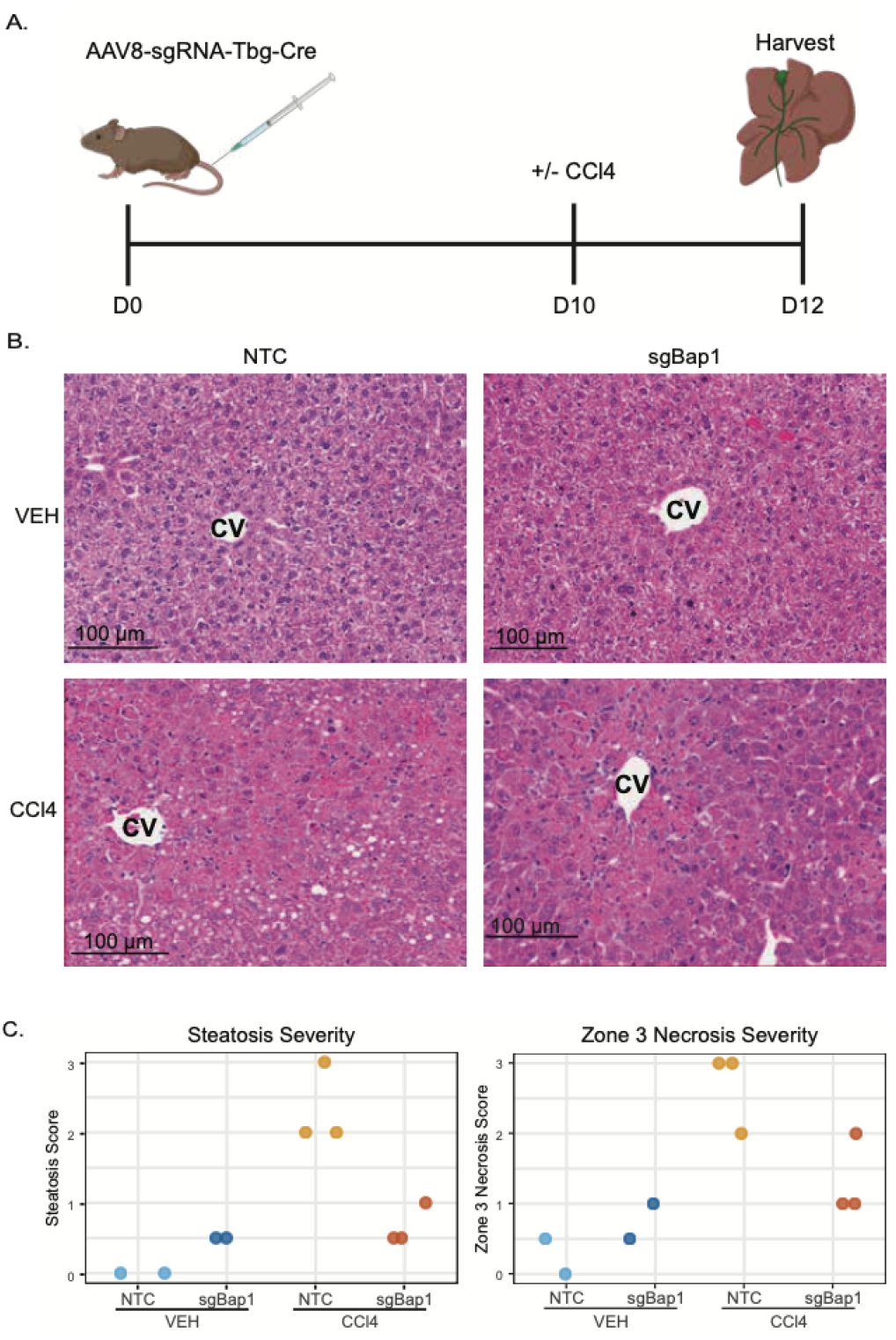
Histologic Changes associated with Bap1 Loss in Hepatocytes. A) Schematic of experimental conditions and timeline. B) Representative hematoxylin and eosin (H&E)-stained images of all experimental conditions. NTC, non-targeting control; VEH, vehicle treated; CCl_4_, carbon tetrachloride treated; CV, central vein. C) Steatosis and zone 3 necrosis grading. Images were graded by increasing severity from 0, +/-(.5), + (1), ++ (2), to +++ (3).

Histologic assessment of liver biopsy sections was performed by a board-certified clinical pathologist (Figure 1B,C). Steatosis was absent in normal NTC livers and upon CCl_4_ treatment was elevated to grade 2 or grade 3. In sgBap1 livers, cells appeared swollen. However, upon CCl_4_ treatment, steatosis was not elevated and few cells displayed fat droplets. Evaluation of zone 3 necrosis showed a similar patterning. Normal NTC livers upon CCl_4_ treatment displayed large necrotic areas with dead hepatocytes that had lost their nuclear staining and had recruitment of inflammatory cells. Vehicle treated sgBAP1 livers had evidence of injury but without cells being completely necrotic yet but after CCl_4_ treatment, damaged areas were not elevated to the same extent as observed in the NTC livers.

### Transcriptional Effect of Bap1 Loss

We recently developed a gene expression score that serves as a proxy for Bap1 activity ^8^. We calculated Bap1 activity scores in the mouse livers to determine the impact of sgBAP1 and liver damage on Bap1 activity (Figure 2A). Both sgBAP1 livers and CCl_4_ treated livers had lower Bap1 activity scores. A similar decrease in BAP1 activity was observed in human liver fibrosis as fibrosis stage worsens ^47^ (Figure 2B). CCl_4_ elicited a strong transcriptional response (n=8,721 genes) with increase in immune and cell cycle signatures and decrease in metabolism gene signature (Supplemental Table 3). Knockdown of Bap1 was associated with 551 genes differentially expressed compared to NTC of which 80% (442 genes) were similarly altered in response to CCl_4_ in vehicle treated livers, suggesting a common pathway impacted by BAP1 loss and liver damage (Figure 2C). The shared gene set had strong upregulation of immune-related pathways from Hallmark and C5 gene sets from MSigDB ^31^ (Figure 2D, E, Supplementary Table 4). GSEA of the differentially expressed genes in Veh-sgBap1 vs Veh-NTC fold changes revealed that 57% of all significantly enriched C5 pathways were immune pathway related (Figure 2F). Taken together, these data demonstrate strong immune cell expression signatures induced by a hepatocyte-specific sgBap1 knockout that shares similarities to CCl_4_ induced damage.

**Figure 2.**
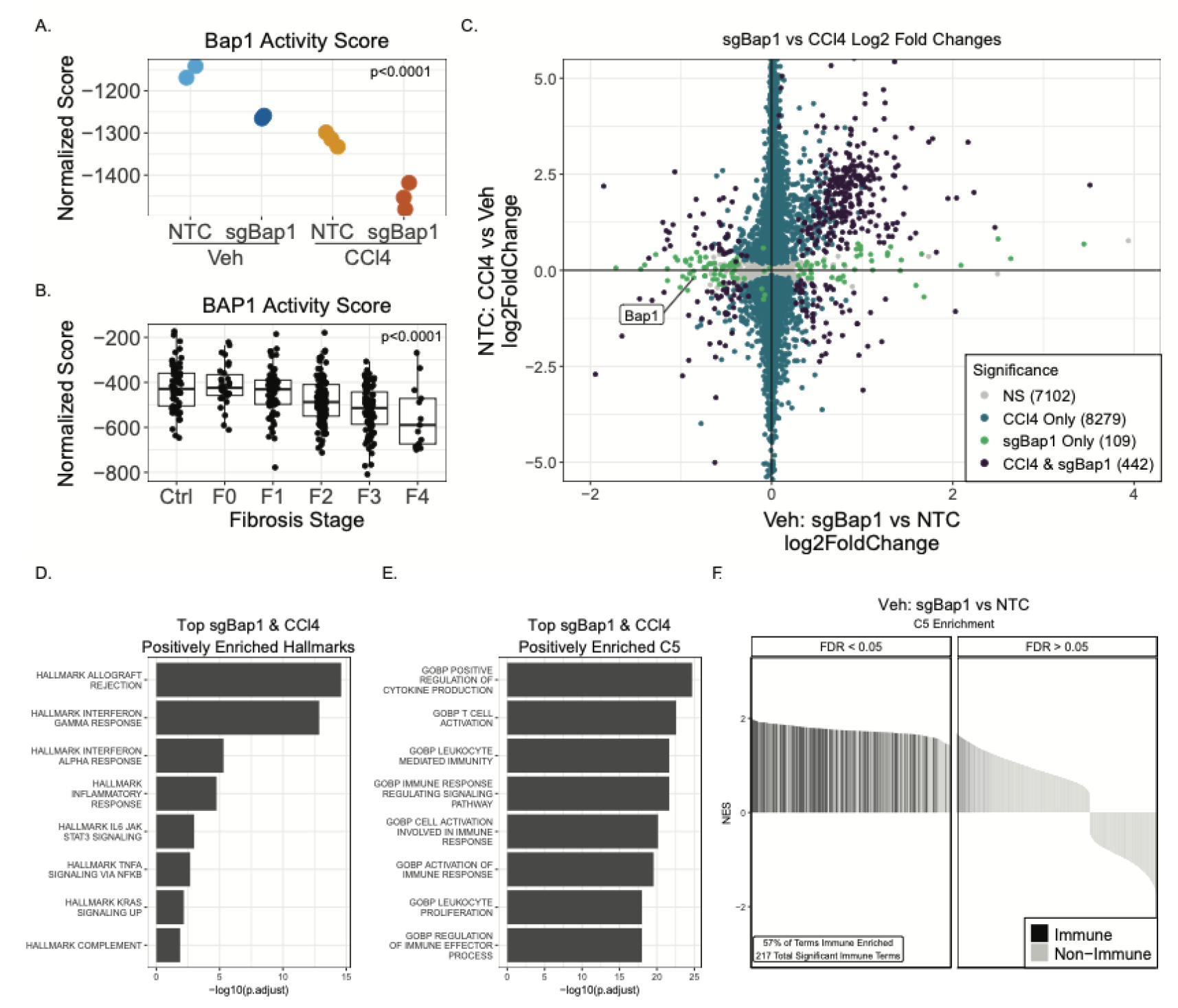
Transcriptional Impact of Bap1 Loss in Liver. A) Bap1 activity score across experimental mouse conditions. Statistical significance assessed by ANOVA test. B) Bap 1 activity score in a human fibrosis data set (ANOVA, p < .0001). C) Scatterplot of sgBap1-altered (Vehicle: sgBap1 vs NTC) and damage-altered (NTC: CCl_4_ vs Vehicle) log2 fold gene expression changes. Points are colored by significance level from DEseq2 (padj < 0.5). Gene set enrichment analysis of shared sgBAP1 and CCl_4_ induced gene expression changes for D) Hallmarks and E) C5 gene sets. The top 8 enriched gene sets are shown ranked by –log10 (adjusted p-value). F) Waterfall plot of normalized enrichment scores (NES) from C5 GSEA performed n knockout specific (Veh: sgBap1 vs NTC) fold changes. Gene sets on the x-axis are ranked by decreasing NES. Black bars indicate gene sets containing immune related keywords.

### BAP1 loss alters the response to damage

We evaluated how Bap1 knockout modulated response to liver damage. We identified 408 genes with a significant interaction between Bap1 loss and CCl_4_ treatment. Unsupervised clustering of these genes revealed four clusters with distinct patterns of expression (Figure 3A).

**Figure 3.**
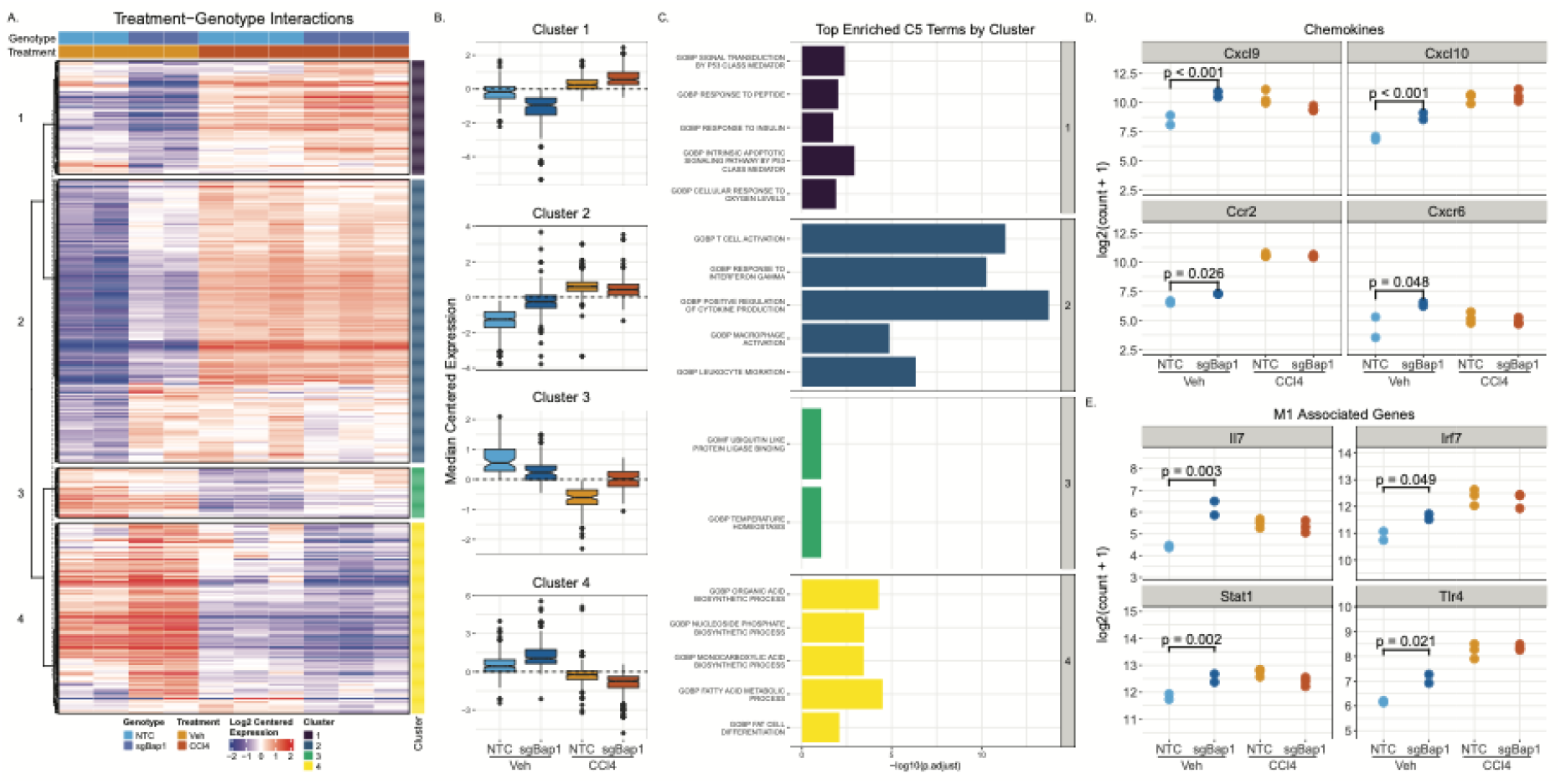
Bap1 Alters Transcriptional Response to Damage. A) Heatmap of median centered normalized log-expression of genes with significant (p < 0.05) treatment-genotype interaction effects. Rows were hierarchically clustered using Pearson distance and average linkage and split into 4 clusters. B) Boxplots of median centered log-expression for genes in clusters identified in A. C) Top enriched MSigDB C5 ontology terms from clusters identified in A plotted by −log10 (adjusted p-value). D) Log2 normalized expression of chemokine and chemokine receptors. E) Log2 normalized expression of genes associated with M1 macrophage polarization.

Cluster 1 patterns showed genes with an overall upregulation of expression with CCl_4_ treatment but were markedly lower in sgBap1 alone livers. (Figure 3B). Cluster 1 was strongly enriched for genes in signal transduction, cell cycle regulation, and stress responses including genes such as Cdkn1a, Egr1, Mdm2, Myc.

Cluster 2 was defined by an immune activated expression phenotype in Veh-sgBap1 that was elevated compared to NTC but lower than the levels observed with CCl_4_ treatment. We observed marked enrichment for signatures related to cytokine signaling, adaptive and innate immune responses, interferon gamma pathway, and tumor necrosis factor pathway (Figure 3C, Supplemental Table 5). We identified several chemokines associated with immune recruitment in the liver (Figure 3D). For example, Cxcl9 and Cxcl10 are expressed by hepatocytes and recruit T cells^49,50^. Ccr2 is a receptor associated with M1 polarized macrophages and monocyte infiltration and Cxcr6 is a natural killer T cell receptor that responds to signals from Kupffer and sinusoidal cells ^49,50^. Vehicle treated sgBap1 livers had elevated macrophage expression patterns characteristic of a pro-inflammatory M1 macrophage phenotype. Il7, Irf7, Stat1, and Tlr4 were upregulated to levels comparable with those observed in CCl_4_ treatment (Figure 3E). However, there was no similar increase in anti-inflammatory M2 macrophage genes Fn1, Msr1, Tgfbr2, or Stat6 (Supplemental Figure 2). These data suggest that M1 macrophages become activated and potentiate a pro-inflammatory environment in livers upon BAP1 loss in hepatocytes.

Cluster 3 was enriched by a small number of transcriptional regulators, including the critical hepatocyte transcriptional regulators Cebpb and Foxo1, which play a role in lipid and glucose metabolism ^51,52^. The expression level of these genes were reduced with damage and sgBap1 relative to the NTC. Taken with cluster 1, these results suggest that Bap1 loss impairs hepatocyte maintenance and transcriptional regulation similar to observed changes with liver damage Cluster 4 was characterized by genes with higher expression in vehicle treated sgBap1 mice. This cluster was enriched with terms related to fatty acid, nucleoside, and organic acid metabolism. These terms were defined by genes known to regulate and alter the lipid profile of the liver. One example, Srebf1, a transcription factor that controls hepatic lipogenesis, is upregulated in the context of NAFLD/MASLD ^53,54^. Furthermore, downstream targets of Srebf1 such as Acly and Elovl6 were upregulated, both of which are involved in lipogenesis (Supplementary Table 3)^55^.

Results from our gene cluster analysis suggest that Bap1 loss alters the cell type composition of the liver. We hypothesized that loss of cell cycle control and upregulation of lipogenesis in hepatocytes due to Bap1 loss induced a proinflammatory environment.

### BAP1 loss alters the cell composition of the liver

To further characterize the changes in cell type composition we performed subcellular spatial transcriptomics on our livers for 100 genes associated with hepatic cell types, differentiation, and proliferation (Figure 4A, Supplemental Table 8). After cell segmentation, we annotated cell types using a liver cell atlas ^45^. Of the 17 cell types in the atlas, we were able to confidently call 8 cell types with at least 100 cells per type (Figure 4B). sgBap1 livers had an increase in Kupffer cells, conventional dendritic cells (cDCs), and T-cells (Figure 4B). Following damage, the increase of Kupffer cells, monocyte-derived cells, cDCs, and T-cells observed in NTC livers was blunted by sgBap1. The cell type labels were applied back to the xy coordinates to evaluate spatial patterns (Figure 4C). The increase in immune cells observed in sgBap1 maintained a general distributed pattern. However, upon damage, sgBap1 did not show the same influx of immune cells to damaged areas.

**Figure 4.**
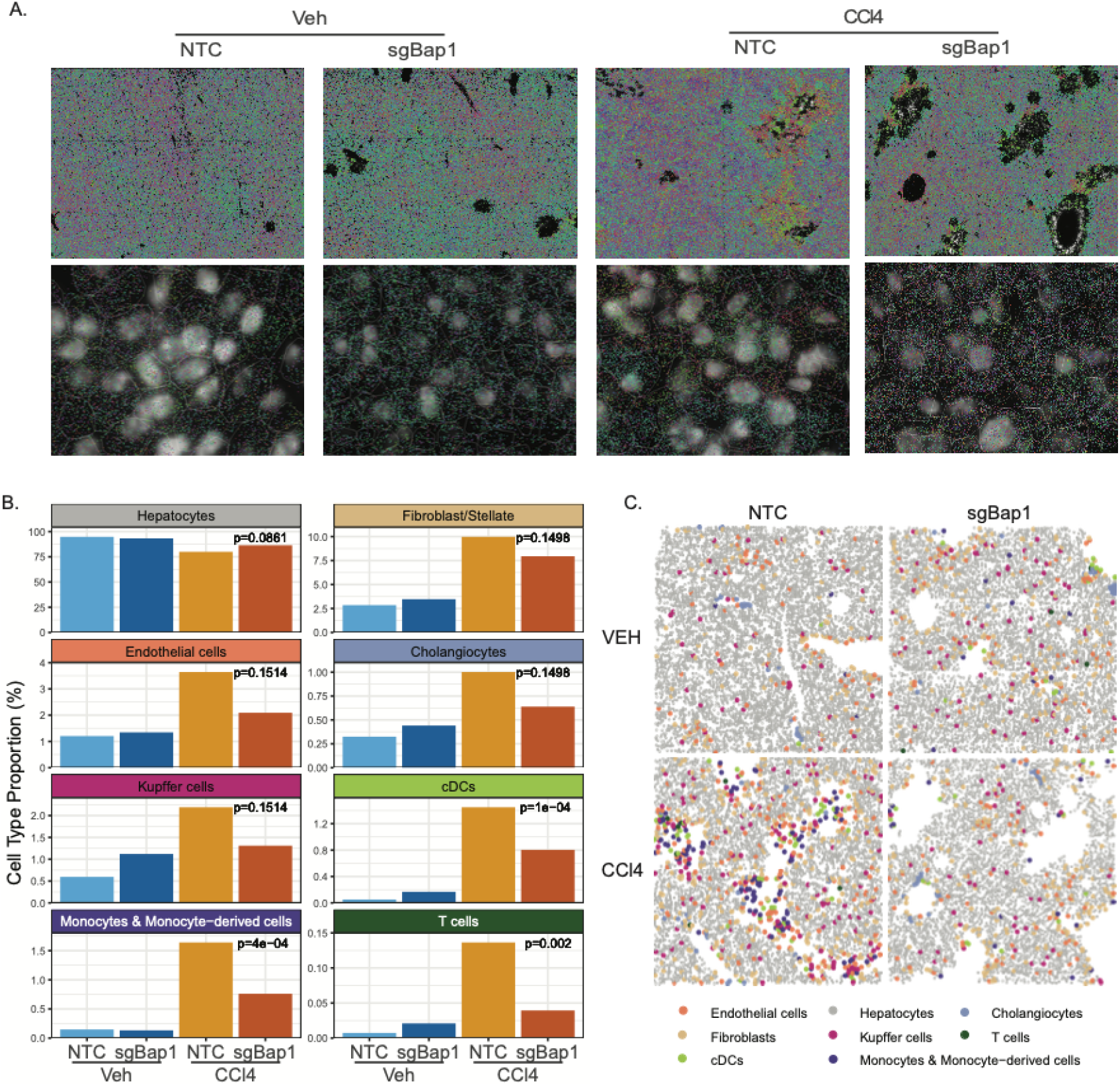
Bap1 Loss in Hepatocytes Alters Cell Populations and Recruitment. A) Representative images from Resolve Biosciences spatial transcriptomics for all experimental conditions. Top, visualization of 100 gene probes; Bottom, cell segmentation overlayed on DAPI stained images with gene probes. B) Proportion of annotated cell types from spatial transcriptomics. Variation in cell populations was assessed by ANOVA. C) Representative reimaged spatial patterning of segmented cells colored by annotated cell type

The altered macrophage signatures identified in the RNA-seq data were further validated by immunohistochemical staining of liver sections for the tissue resident macrophages. We observed similar trends, showing an increased staining and density of F4/80 in sgBap1 livers but still retained a dispersed spatial patterning as in the NTC livers (Figure 5A). After damage, NTC livers drastically increased staining for F4/80 and the cells clustered near damage around central veins with staining appearing more diffuse (Figure 5A). However, in sgBap1 livers the response to damage was less robust with less intense staining and reduced recruitment to areas of damage. The visual changes were confirmed with the semi-quantitative H-score, which was increased in sgBap1 livers with Veh, but displayed a muted and altered response with CCl_4_ treatment (Figure 5B). These data were consistent with bulk RNA-seq expression levels for Adgre1, the gene encoding F4/80 (Figure 5C).

**Figure 5.**
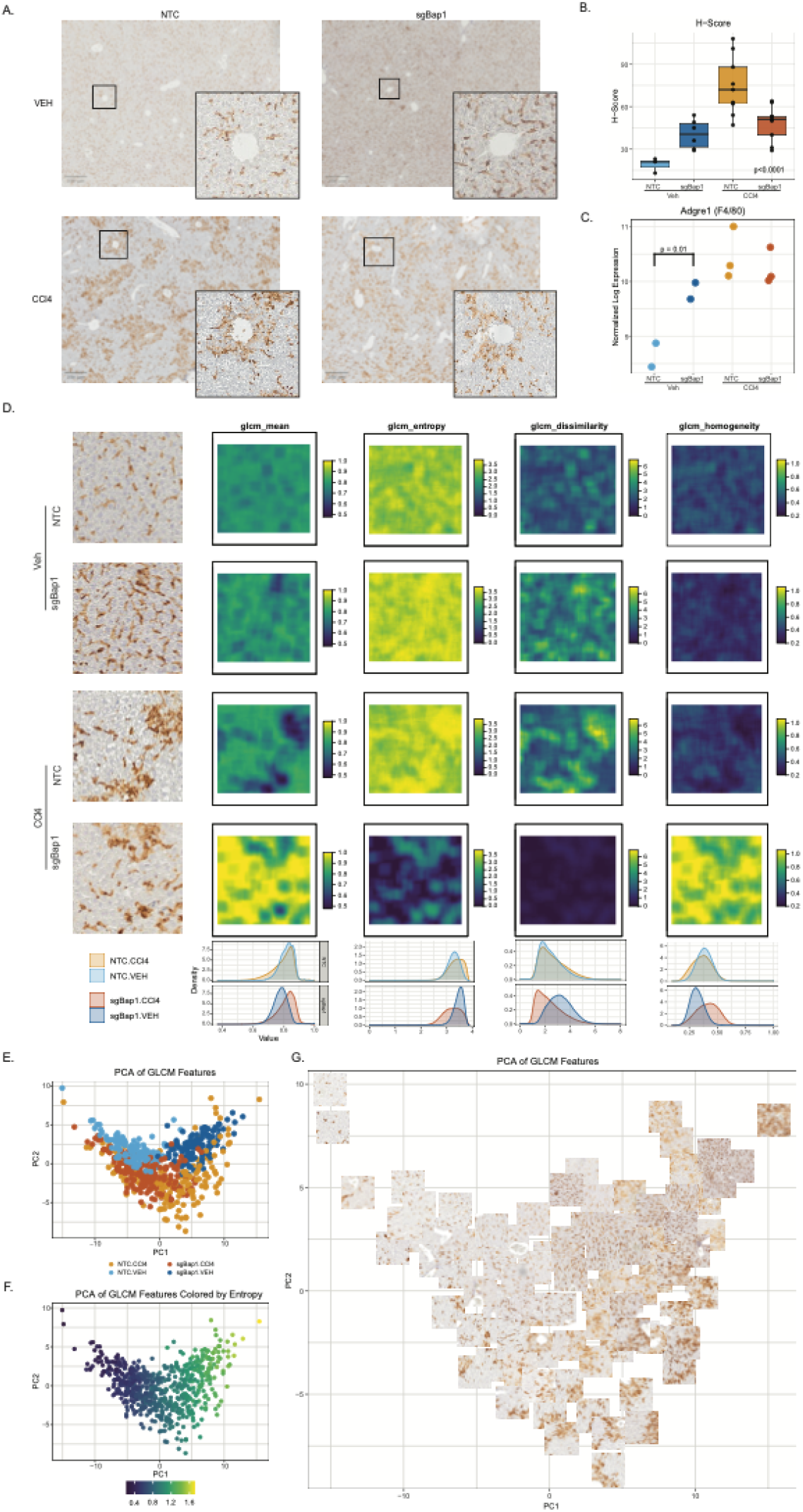
Altered macrophage spatial patterning with Bap1 loss. A) Representative F4/80-stained images of all experimental conditions. Image inlay of magnified area around a central vein. B) H-score of F4/80 immunohistochemical stain. Statistical significance is assessed with one way ANOVA. C) Log2 normalized gene expression of Adgre1 (F4/80). D) Image-based texture analysis using grey level co-occurrence matrix analysis (GLCM) on F4/80 IHC images. Representative tiles of all 4 experimental conditions for four of 7 GLCM features – mean, entropy, dissimilarity, and homogeneity (Supplemental Figure 3 for the other four features). Bottom, distributions of all pixel values for each filter colored by condition. Principal component analysis (PCA) of tile level GLCM summary statistics colored by E) condition, F) GLCM entropy value, or G) subset of Tile-level representations.

Given the changes in macrophage distribution that we observed, we sought to quantitatively assess changes in tissue structure and spatial organization of macrophages using texture analysis ^434^ This approach allowed us to statistically quantify and summarize the distribution, amount, and size of cells in F4/80 stained IHC images. Each image was broken into small tiles and analyzed for 7 texture features (Supplemental Figure 3) at a pixel level and summarized at a tile level. These features captured variances both with sgBap1 and CCl_4_ treatment (Figure 5D, Supplemental Figure 4). NTC liver tiles had high entropy and variance, features defining the dispersion of F4/80 stain cells across tissues. In sgBap1, we noted increased entropy, dissimilarity, and contrast and decreased homogeneity and variance suggesting a shift towards a more complex, heterogeneous, and less uniform texture. With damage, sgBap1 tissues showed dramatic differences in texture patterns for features. We plotted summary metrics of each feature for each tile using PCA and found separation of image tiles by both sgBap1 and CCl_4_ status (Figure 5E). The PCA loadings provided insights into key features driving the clustering (Supplemental Figure 5, Supplemental Table 11). The first PC was driven by features indicative of cell number (Figure 5F), while PC2 was driven by features of variation (Supplemental Figure 6). For a more interpretable visualization, we plotted a subset of the points using their tile image (Figure 5G). While the sgBap1 tiles show clear increase in F4/80 stain similar to CCl_4_ treated samples, upon damage sgBap1 tissues show altered macrophage recruitment with lower diffuseness and recruitment to central veins. Together, these data suggest that a modest increase in macrophages occurs in undamaged sgBap1 livers, but that macrophage recruitment to damage is altered by Bap1 loss in hepatocytes. Moreover, this analysis highlights the impact of quantitative assessment of spatial patterns which can reveal more detailed insights into how loss of Bap1 impacts the spatial distribution of macrophages.

## Discussion

Germline loss of BAP1 leads to a tumor predisposition syndrome and somatic mutations of BAP1 are common in primary liver cancer, uveal melanoma, and mesothelioma^56–58^. However, few studies have interrogated the function of BAP1 outside of cancer. We sought to define the role of BAP1 in normal liver cells and to determine if these changes contribute to a role for Bap1 as a potential early driver for liver disease and later cancer risk. The activation of hepatic stellate cells by damaged hepatocytes is a key initial driver of the liver damage response, and dysfunction of this interaction can lead to maladaptive repair and fibrosis ^59,60^. Therefore, we investigated Bap1’s role in normal hepatocytes and how it contributes in response to acute liver damage. We deleted Bap1 specifically in hepatocytes which led to increased inflammation and altered recruitment of immune cells in the uninjured liver.

These data suggest that Bap1 on its own activates an inflammatory response that mimics changes seen after liver injury. Notably, similar ‘pre-injured’ states were found in the histology data with changes to macrophage localization and steatosis. The link between Bap1 loss and an immune response has been observed in several prior studies. However, Bap1 loss does not lead to a consistent directionality of that immune response across different tissues. For example, in uveal melanoma loss of Bap1 leads to an immunosuppressive environment by disrupting NF-kB signaling, while in mesothelioma Bap1 loss is associated with interferon alpha and gamma and in increase inflammatory response ^18,61^. Our data support Bap1 loss as an inducer of an inflammatory response and many of these inflammatory signatures were also upregulated in normal livers in response to damage. We noted an increase in chemokine expression in sgBap1 livers which are likely acting as recruiters for the increased immune cells we observed. While F4/80 is not a Kupffer cell specific marker, our gene expression data showed an increase in Clec4f a Kupffer cell specific marker in the mouse liver suggesting the increase is more dependent on resident macrophages than infiltrating macrophages. There was also increased expression of genes involved in the inflammasome potentially reflecting a primed proinflammatory milieu driven by Bap1 loss in hepatocytes.

Paradoxically, the transcriptional and immune cell response to damage of Bap1-depleted livers was blunted compared to wild-type livers. We noted this reduced immune response in both the expression data and in the histology data. CCl_4_ treated sgBap1 livers had reduced zone 3 necrosis and reduced migration of F4/80 macrophages to damage at the central vein. Recent work has shown that during damage zone 3 necrotic regions are predominantly populated and cleared by infiltrating monocyte-derived macrophages while Kupffer Cells predominately stay in zones 1 and 2 ^62^. Even though inflammatory pathways were already partially activated with Bap1 loss, there is a disruption of signaling of immune response to damage. This could be due to changes in how cytokine signals are sent or received by adjacent cells or could reflect an exhaustion of the liver environment driven by the premature activation of an immune response in Bap1-depleted hepatocytes.

While we do not know the direct mechanism through which hepatocytes influence immune cell recruitment and response, the increase in expression of chemokines in uninjured Bap1 depleted livers that does not become further elevated with damage, suggests a dysfunction cross talk mechanism. Together, these experiments suggest an early role in priming the liver for maladaptive repair and support Bap1 loss as an early event that contributes to later liver disease; an idea supported by human fibrosis data where Bap1 activity was reduced with increasing severity of fibrosis. Our experiments implicate Bap1 having a broader role in maintaining homeostasis in the uninjured liver. Future studies will need to carefully dissect the temporal roles of Bap1 and its molecular functions to maintain proper liver function and response to injury.

## References

1. Wang, Z., Liu, Z., Lv, M. et al. Novel histone modifications and liver cancer: emerging frontiers in epigenetic regulation. Clin Epigenet 17, 30 (2025). 10.1186/s13148-025-01838-8

2. Zhao, S., Allis, C.D. & Wang, G.G. The language of chromatin modification in human cancers. Nat Rev Cancer 21, 413–430 (2021). 10.1038/s41568-021-00357-x

3. Flavahan, W. A., Gaskell, E. & Bernstein, B. E. Epigenetic plasticity and the hallmarks of cancer. Science 357, (2017).

4. Hodges, C., Kirkland, J. G. & Crabtree, G. R. The Many Roles of BAF (mSWI/SNF) and PBAF Complexes in Cancer. Cold Spring Harb. Perspect. Med. 6, (2016).

5. Blackledge, N. P. et al. Variant PRC1 complex-dependent H2A ubiquitylation drives PRC2 recruitment and polycomb domain formation. Cell 157, 1445–1459 (2014).

6. Grindheim, J. M., Nicetto, D., Donahue, G. & Zaret, K. S. Polycomb Repressive Complex 2 Proteins EZH1 and EZH2 Regulate Timing of Postnatal Hepatocyte Maturation and Fibrosis by Repressing Genes With Euchromatic Promoters in Mice. Gastroenterology 156, 1834–1848 (2019).

7. Yan, Q. et al. RNF2 Mediates Hepatic Stellate Cells Activation by Regulating ERK/p38 Signaling Pathway in LX-2 Cells. Front Cell Dev Biol 9, 634902 (2021).

8. Sturgill, I. R., Raab, J. R. & Hoadley, K. A. Expanded detection and impact of BAP1 alterations in cancer. NAR Cancer 6, zcae045 (2024).

9. Cancer Genome Atlas Research Network. Electronic address: & Cancer Genome Atlas Research Network. Comprehensive and Integrative Genomic Characterization of Hepatocellular Carcinoma. Cell 169, 1327–1341.e23 (2017).

10. Farshidfar, F. et al. Integrative Genomic Analysis of Cholangiocarcinoma Identifies Distinct IDH-Mutant Molecular Profiles. Cell Rep. 18, 2780–2794 (2017).

11. Damrauer, J. S. et al. Genomic characterization of rare molecular subclasses of hepatocellular carcinoma. Commun Biol 4, 1150 (2021).

12. Baughman, J.M., Rose, C.M., Kolumam G., Webster, J.D., Wilkerson, E.M., Merrill, A.E., Rhoads, T.W., Noubade, R., Katavolos, P., Lesch, J., et al. (2016) NeuCode Proteomics Reveals Bap1 Regulation of Metabolism. Cell Rep., 16, 583–595.

13. Artegiani, B. et al. Probing the Tumor Suppressor Function of BAP1 in CRISPR-Engineered Human Liver Organoids. Cell Stem Cell 24, 927–943.e6 (2019).

14. Zhang, T. et al. An epigenetic pathway regulates MHC-II expression and function in B cell lymphoma models. J. Clin. Invest. 135, (2025).

15. Liang, Y. et al. B-cell intrinsic regulation of antibody mediated immunity by histone H2A deubiquitinase BAP1. Front. Immunol. 15, 1353138 (2024).

16. Radhakrishnan, D. et al. Deubiquitinase BAP1 is crucial for surface expression of T cell receptor (TCR) complex, T cell-B cell conjugate formation, and T cell activation. J. Leukoc. Biol. 117, (2024).

17. Chang, H. et al. Loss of histone deubiquitinase Bap1 triggers anti-tumor immunity. Cell. Oncol. (2024) doi:10.1007/s13402-024-00978-y.

18. Zhang, C. & Wu, S. BAP1 mutations inhibit the NF-κB signaling pathway to induce an immunosuppressive microenvironment in uveal melanoma. Mol. Med. 29, 126 (2023).

19. Camp, S. Y. et al. Single-cell epigenetic profiling reveals an interferon response-high program associated with BAP1 deficiency in kidney cancer. bioRxivorg (2024) doi:10.1101/2024.11.15.623837.

20. Bankhead, P., Loughrey, M.B., Fernández, J.A. et al. QuPath: Open source software for digital pathology image analysis. Sci Rep 7, 16878 (2017). 10.1038/s41598-017-17204-5

21. Wickham, H., Chang, W., Henry, L., Pedersen, T. L., Takahashi, K., Wilke, C., Woo, K., Yutani, H., Dunnington, D., Brand, T. van den, Posit, & PBC. (2025). ggplot2: Create Elegant Data Visualisations Using the Grammar of Graphics (Version 4.0.0)

22. Dobin, Alexander et al. “STAR: ultrafast universal RNA-seq aligner.” Bioinformatics (Oxford, England) vol. 29,1 (2013): 15–21. doi:10.1093/bioinformatics/bts635

23. Patro, Rob et al. “Salmon provides fast and bias-aware quantification of transcript expression.” Nature methods vol. 14,4 (2017): 417–419. doi:10.1038/nmeth.4197

24. Li, Heng, et al. “The Sequence Alignment/Map Format and SAMtools.” Bioinformatics, vol. 25, no. 16, Aug. 2009, pp. 2078–79. Silverchair, 10.1093/bioinformatics/btp352.

25. Andrews, S. (2010) FastQC A Quality Control Tool for High Throughput Sequence Data. - References - Scientific Research Publishing. https://www.scirp.org/reference/referencespapers?referenceid=2781642. Accessed 3 Oct. 2025.

26. Love, Michael I., et al. “Moderated Estimation of Fold Change and Dispersion for RNA-Seq Data with DESeq2.” Genome Biology, vol. 15, no. 12, Dec. 2014, p. 550. BioMed Central, 10.1186/s13059-014-0550-8.

27. Zhu, Anqi, et al. “Heavy-Tailed Prior Distributions for Sequence Count Data: Removing the Noise and Preserving Large Differences.” Bioinformatics, vol. 35, no. 12, June 2019, pp. 2084–92. Silverchair, 10.1093/bioinformatics/bty895.

28. Gu, Zuguang. “Complex Heatmap Visualization.” iMeta, vol. 1, no. 3, 2022, p. e43. Wiley Online Library, 10.1002/imt2.43.

29. Liberzon, Arthur, et al. “The Molecular Signatures Database Hallmark Gene Set Collection.” Cell Systems, vol. 1, no. 6, Dec. 2015, pp. 417–25. ScienceDirect, 10.1016/j.cels.2015.12.004.

30. Subramanian, Aravind, et al. “Gene Set Enrichment Analysis: A Knowledge-Based Approach for Interpreting Genome-Wide Expression Profiles.” Proceedings of the National Academy of Sciences, vol. 102, no. 43, Oct. 2005, pp. 15545–50. pnas.org (Atypon), 10.1073/pnas.0506580102.

31. Castanza, Anthony S., et al. “Extending Support for Mouse Data in the Molecular Signatures Database (MSigDB).” Nature Methods, vol. 20, no. 11, Nov. 2023, pp. 1619–20. http://www.nature.com, 10.1038/s41592-023-02014-7.

32. CRAN: Package Msigdbr. https://cran.r-project.org/web/packages/msigdbr/index.html. Accessed 3 Oct. 2025.

33. Wu, Tianzhi, et al. “clusterProfiler 4.0: A Universal Enrichment Tool for Interpreting Omics Data.” The Innovation, vol. 2, no. 3, Aug. 2021, p. 100141. ScienceDirect, 10.1016/j.xinn.2021.100141.

34. Yu, Guangchuang, et al. “clusterProfiler: An R Package for Comparing Biological Themes Among Gene Clusters.” OMICS: A Journal of Integrative Biology, vol. 16, no. 5, May 2012, pp. 284–87. liebertpub.com (Atypon), 10.1089/omi.2011.0118.

35. Petukhov, V., Xu, R.J., Soldatov, R.A. et al. Cell segmentation in imaging-based spatial transcriptomics. Nat Biotechnol 40, 345–354 (2022). 10.1038/s41587-021-01044-w

36. Cable, Dylan M., et al. “Robust Decomposition of Cell Type Mixtures in Spatial Transcriptomics.” Nature Biotechnology, vol. 40, no. 4, Apr. 2022, pp. 517–26. http://www.nature.com, 10.1038/s41587-021-00830-w.

37. Guilliams, Martin et al. “Spatial proteogenomics reveals distinct and evolutionarily conserved hepatic macrophage niches.” Cell vol. 185,2 (2022): 379-396.e38. doi:10.1016/j.cell.2021.12.018

38. Phipson, Belinda et al. “propeller: testing for differences in cell type proportions in single cell data.” Bioinformatics (Oxford, England) vol. 38,20 (2022): 4720–4726. doi:10.1093/bioinformatics/btac582

39. Mouselimis, Lampros, et al. OpenImageR: An Image Processing Toolkit. Version 1.3.0, 8 July 2023. R-Packages, https://cran.r-project.org/web/packages/OpenImageR/index.html.

40. Pau, Grégoire, et al. “EBImage—an R Package for Image Processing with Applications to Cellular Phenotypes.” Bioinformatics, vol. 26, no. 7, Apr. 2010, pp. 979–81. Silverchair, 10.1093/bioinformatics/btq046

41. CRAN: Package Raster. https://cran.r-project.org/web/packages/raster/index.html. Accessed 3 Oct. 2025.

42. Hijmans, Robert J., et al. Terra: Spatial Data Analysis. Version 1.8-70, 27 Sept. 2025. R-Packages, https://cran.r-project.org/web/packages/terra/index.html.

43. Haralick, Robert M., et al. “Textural Features for Image Classification.” IEEE Transactions on Systems, Man, and Cybernetics, SMC-3, no. 6, Nov. 1973, pp. 610–21. IEEE Xplore, 10.1109/TSMC.1973.4309314.

44. Zvoleff, Alex. Glcm: Calculate Textures from Grey-Level Co-Occurrence Matrices (GLCMs). Version 1.6.5, 26 Feb. 2020. R-Packages, https://cran.r-project.org/web/packages/glcm/index.html.

45. Meyer [aut, David, et al. E1071: Misc Functions of the Department of Statistics, Probability Theory Group (Formerly: E1071), TU Wien. Version 1.7-16, 16 Sept. 2024. R-Packages, https://cran.r-project.org/web/packages/e1071/index.html.

46. Yu, Guangchuang, and Shuangbin Xu. Ggimage: Use Image in “Ggplot2.” Version 0.3.4, 24 Aug. 2025. R-Packages, https://cran.r-project.org/web/packages/ggimage/index.html.

47. Chen, Tianyi et al. “Hepatocyte Smoothened Activity Controls Susceptibility to Insulin Resistance and Nonalcoholic Fatty Liver Disease.” Cellular and molecular gastroenterology and hepatology vol. 15,4 (2023): 949–970. doi:10.1016/j.jcmgh.2022.12.008

48. Derdak, Zoltan, et al. “Inhibition of P53 Attenuates Steatosis and Liver Injury in a Mouse Model of Non-Alcoholic Fatty Liver Disease.” Journal of Hepatology, vol. 58, no. 4, Apr. 2013, pp. 785–91. PubMed, 10.1016/j.jhep.2012.11.042.

49. Cao, Sheng, et al. “Regulation and Functional Roles of Chemokines in Liver Diseases.” Nature Reviews. Gastroenterology & Hepatology, vol. 18, no. 9, Sept. 2021, pp. 630–47. PubMed, 10.1038/s41575-021-00444-2.

50. Marra, Fabio, and Frank Tacke. “Roles for Chemokines in Liver Disease.” Gastroenterology, vol. 147, no. 3, Sept. 2014, pp. 577-594.e1. PubMed, 10.1053/j.gastro.2014.06.043.

51. Tikhanovich, Irina et al. “Forkhead box class O transcription factors in liver function and disease.” Journal of gastroenterology and hepatology vol. 28 Suppl 1,0 1 (2013): 125–31. doi:10.1111/jgh.12021

52. Wang, Bin, et al. “C/EBPβ Contributes to Hepatocyte Growth Factor-Induced Replication of Rodent Hepatocytes.” Journal of Hepatology, vol. 43, no. 2, Aug. 2005, pp. 294–302. http://www.journal-of-hepatology.eu, 10.1016/j.jhep.2005.02.029.

53. Li, Na, et al. “SREBP Regulation of Lipid Metabolism in Liver Disease, and Therapeutic Strategies.” Biomedicines, vol. 11, no. 12, Dec. 2023, p. 3280. PubMed, 10.3390/biomedicines11123280.

54. Im, Seung-Soon, et al. “Linking Lipid Metabolism to the Innate Immune Response in Macrophages through Sterol Regulatory Element Binding Protein-1a.” Cell Metabolism, vol. 13, no. 5, May 2011, pp. 540–49. PubMed, 10.1016/j.cmet.2011.04.001.

55. Horton, Jay D et al. “Combined analysis of oligonucleotide microarray data from transgenic and knockout mice identifies direct SREBP target genes.” Proceedings of the National Academy of Sciences of the United States of America vol. 100,21 (2003): 12027–32. doi:10.1073/pnas.1534923100

56. Rai, K et al. “Comprehensive review of BAP1 tumor predisposition syndrome with report of two new cases.” Clinical genetics vol. 89,3 (2016): 285–94. doi:10.1111/cge.12630

57. Harbour, J William et al. “Frequent mutation of BAP1 in metastasizing uveal melanomas.” Science (New York, N.Y.) vol. 330,6009 (2010): 1410–3. doi:10.1126/science.1194472

58. Cheung, Mitchell, and Joseph R Testa. “BAP1, a tumor suppressor gene driving malignant mesothelioma.” Translational lung cancer research vol. 6,3 (2017): 270–278. doi:10.21037/tlcr.2017.05.03

59. Seki, E. & Brenner, D. A. Recent advancement of molecular mechanisms of liver fibrosis. J. Hepatobiliary Pancreat. Sci. 22, 512–518 (2015).

60. Kamm, Dakota R, and Kyle S McCommis. “Hepatic stellate cells in physiology and pathology.” The Journal of physiology vol. 600,8: 1825–1837 (2022).

61. Figueiredo, Carlos R et al. “Loss of BAP1 expression is associated with an immunosuppressive microenvironment in uveal melanoma, with implications for immunotherapy development.” The Journal of pathology vol. 250,4 (2020): 420–439. doi:10.1002/path.5384

62. Feng, Dechun et al. “Characterisation of macrophages in healthy and diseased livers in mice: identification of necrotic lesion-associated macrophages.” eGastroenterology vol. 3,2 e100189. 7 Apr. 2025, doi:10.1136/egastro-2025-100189

