## Supplemental Figures for "BAP1 deletion in hepatocytes primes an inflammatory transcriptional response"

**Supplemental Figures and Table Descriptions**

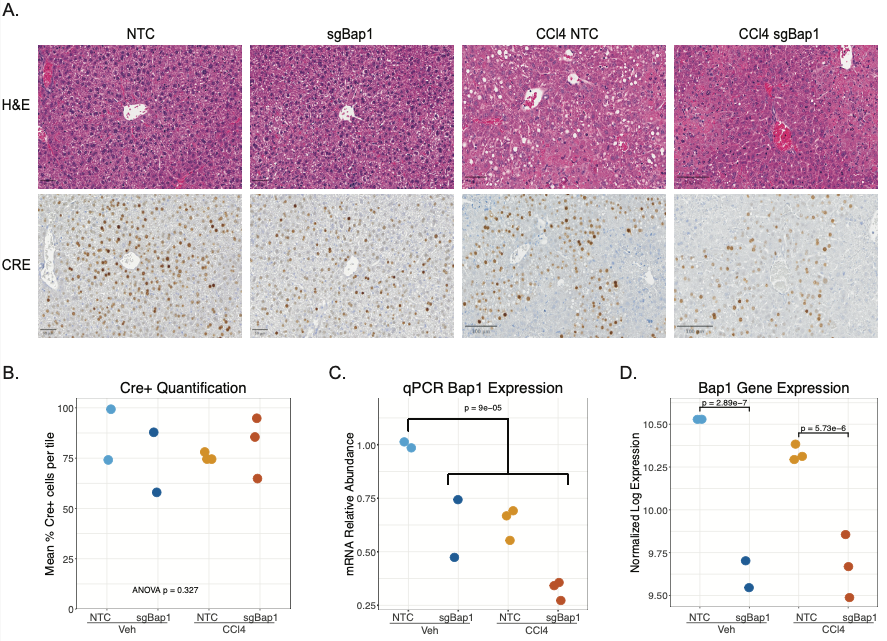

**Supplemental Figure 1. Histologic and Expression quantification of Cre and Bap1.** A) Representative Hematoxylin and eosin (H&E)-stained and immunohistochemistry stained Cre images. B) qPCR of Cre expression levels. C) qPCR of Bap1 expression levels. D) Bulk RNA-seq gene expression levels of Bap1.

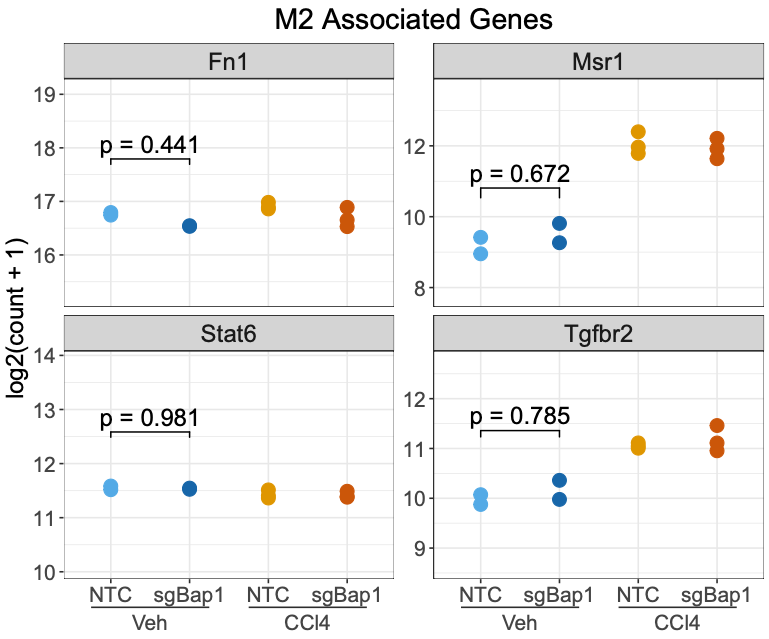

**Supplemental Figure 2. Pro-inflammatory M2 macrophage gene expression.** Log2 normalized expression of genes associated with M2 macrophage polarization.

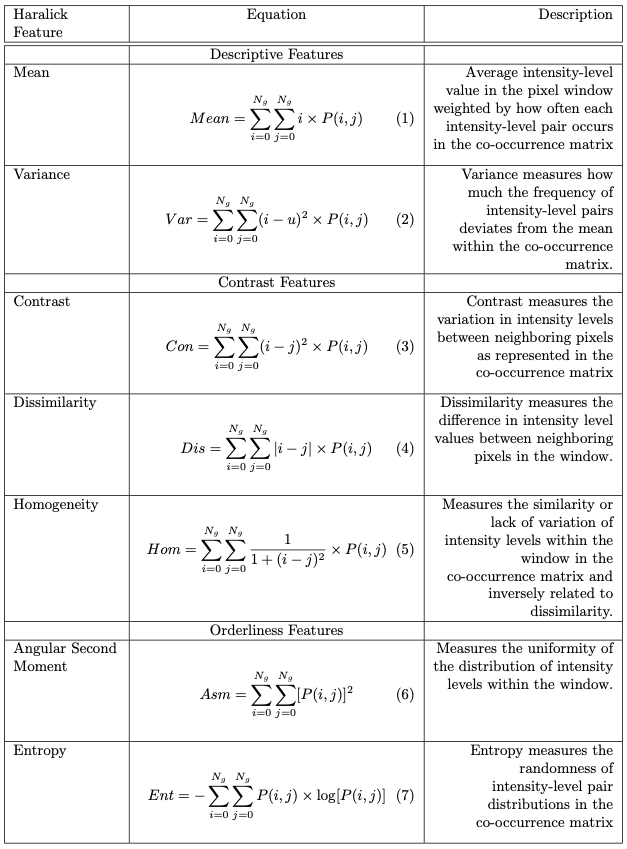

**Supplemental Fig 3. Image-based texture analysis.** Equations and descriptions for Haralick features performed in GLCM texture analysis Fig 5D-G^43,44^.

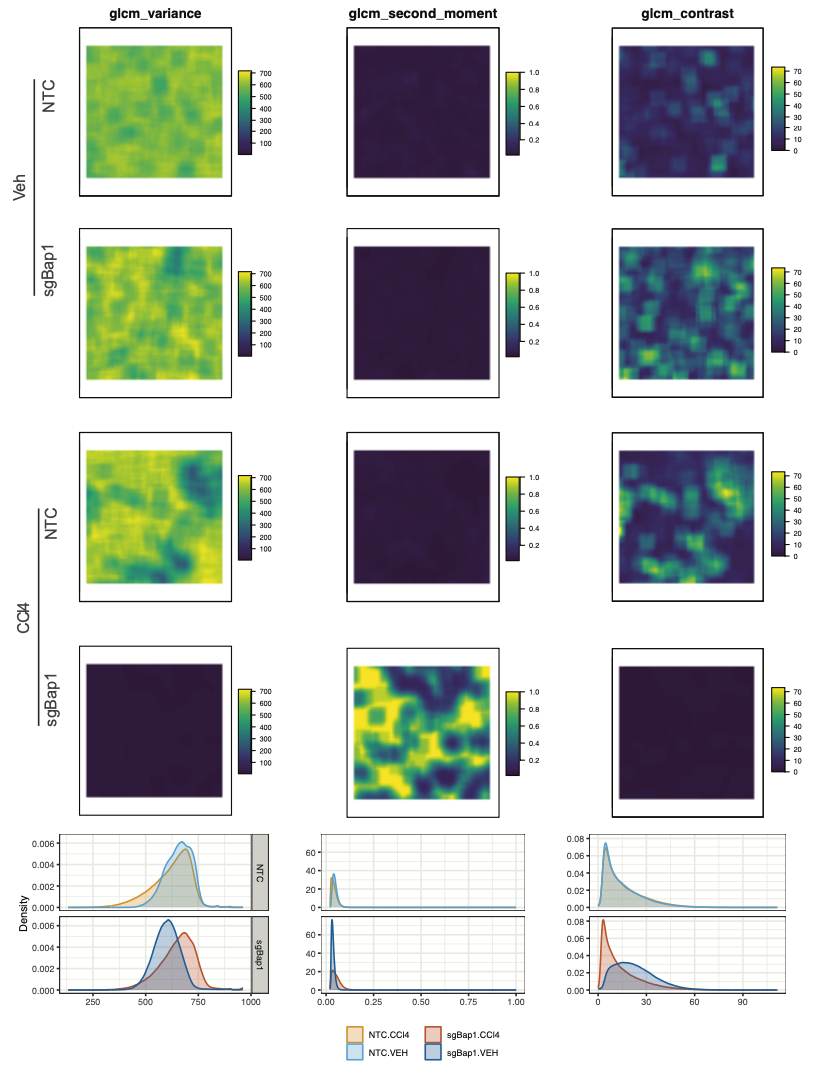

**Supplemental Fig 4. Additional image-based texture analysis using grey level co-occurrence matrix analysis (GLCM) on F4/80 IHC images.** Representative tiles of all experimental conditions for remaining 3 of 7 GLCM features – contrast, variance, and second moment. Bottom, distribution of all pixel values for each filter colored by condition.

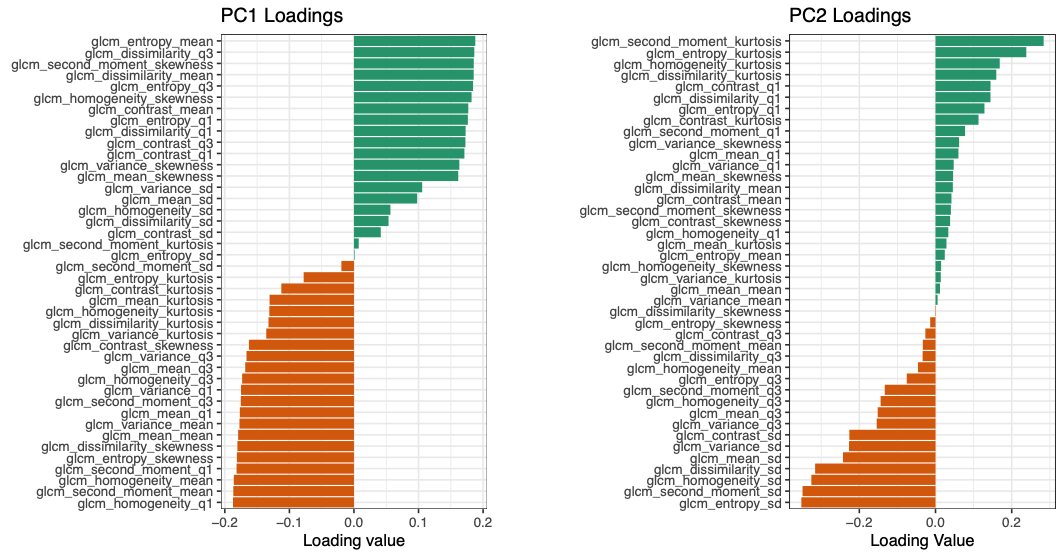

**Supplemental Figure 5. Waterfall plot of PCA loading values for both PC1 and PC2.**

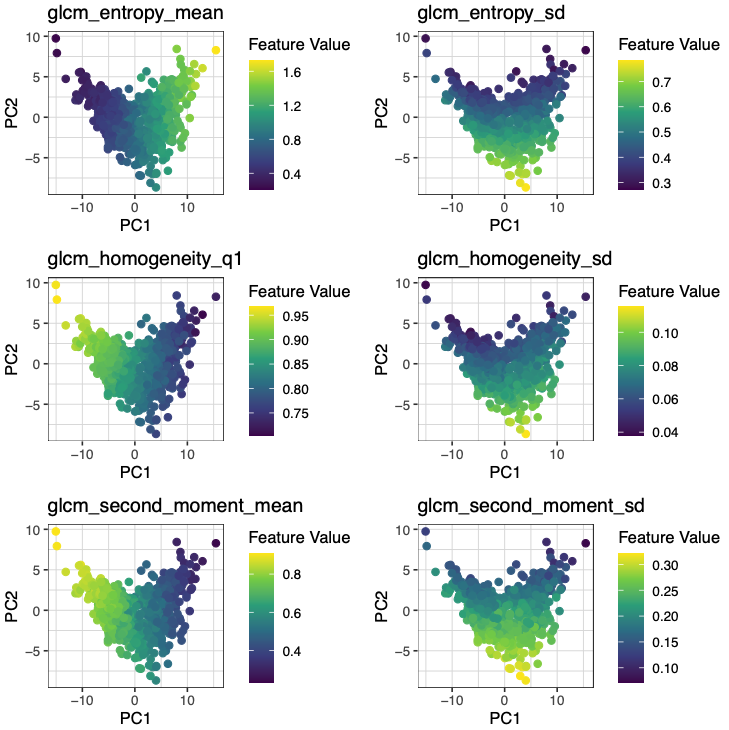

**Supplemental Figure 6. PCA of GLCM texture values colored by top 3 PC loadings from PC1 (left) and PC2 (right).**

**Supplemental Tables**

1. qPCR results for BAP1.
2. Pathological summary of H&E slides for Necrosis and Steatosis related to Figure 1C.
3. Shrunken log2Fold changes of all genes. Each tab is a specific comparison noted below
   a. sgbap1.Veh vs NTC
   b. SgBap1.CCl4 vs NTC
   c. NTC.CCl4 vs VEH
   d. sgBap1.CCl4 vs Veh
4. GSEA of shared significantly enriched genes from Figure 2C. Tab for each analysis noted below.
   a. Hallmark pathways Figure 2D.
   b. C5 pathways Figure 2 E,F.
5. GSEA of genes with significant interaction effects. Tab for each gene set noted below
   a. Interaction genes that had significant change and their cluster in Figure 3A.
   b. C5 gene set enrichment output of clusters from interaction terms Figure 3C.
6. Immune Terms identified from C5 MSigDB data set for Figure 2F.
7. Bap1 activity scores of samples.Tab for each data set noted below.
   a. Bap1 activity score of fibrosis dataset.
   b. Bap1 activity score of hepatic Bap1 KO dataset.
8. Gene Probes Selected for Spatial Transcriptomics in Resolve Biosciences.
9. F4/80 images and their reference patches as well as tissue patches from Figure 5D-G.
10. PC values from GLCM analysis with treatment groupings from Figure 5D-G.
11. PC loading values from GLCM analysis with treatment groupings from Figure 5D-G.
